# HLA variants and TCR diversity against SARS-CoV-2 in the pre-COVID-19 era

**DOI:** 10.1101/2022.09.14.507948

**Authors:** Stéphane Buhler, Zuleika Calderin Sollet, Florence Bettens, Antonia Schaefer, Marc Ansari, Sylvie Ferrari-Lacraz, Jean Villard

## Abstract

HLA antigen presentation and T-cell immunity are critical to control viral infection such as SARS-CoV-2. This study performed on samples collected in the pre-COVID-19 era demonstrates that individuals are fully equiped at the genetic level in terms of TCR repertoire and HLA variants to recognize and kill SARS-CoV-2 infected cells. HLA diversity, heterologous immunity and random somatic TCR recombination could explain these observations.

## Main text

Acute infection caused by the SARS-CoV-2 coronavirus has resulted in a worldwide burden since the start of the COVID-19 pandemic. Disease outcomes vary among infected individuals, from asymptomatic to life-threatening symptoms. Host and genetic factors have been identified that contribute to COVID-19 susceptibility ^1^. T-cell mediated immunity plays a key role and provides a lasting protection in convalescent people ^2^. A trillion of T cells circulate in the human body, each cell harboring a specific T-cell receptor (TCR) generated by quasi-random somatic gene recombination. The TCR is the functional unit allowing T cells to recognize their cognate ligands as complexes formed by a peptide and HLA molecule (pHLA) ^3^. For this system to operate, the T-cell repertoire needs to cover a vast array of peptides. The basis of pHLA-TCR interactions is thus cross-reactive ^4^. In the context of COVID-19, it has been shown that unexposed people can develop significant reactivity against SARS-CoV-2 derived peptides due to cross-reactivity of memory T cells specific to common cold coronaviruses (HCovs) ^5,6^. This heterologous immunity could be a contributing factor in disease outcome. As T cells screen foreign antigens through the lens of highly polymorphic HLA molecules, studies have focused on an association between HLA variation and COVID-19 ^7^. Numerous alleles have been associated, but without clear consensus, while some large studies have failed to confirm an effect ^8,9^.

In this study, we analyzed a cohort of patients having received an allogeneic hematopoietic stem cell transplantation (alloHSCT) before 2019 and their corresponding donors ^10^. High resolution HLA genotyping was performed and the TCR CDR3β repertoire was surveyed by immunosequencing. AlloHSCT is a unique model to analyse the T cell repertoire reconstitution in human and give us the opportunity to explore the presence of SARS-CoV2 clonotypes in unexposed subjects before alloHSCT and in the context of immune reconstitution one year after alloHSCT ^11^. Among the 3,565,567 clonotypes analysed, a total of 171,203 shared their CDR3β sequence with a SARS-CoV-2 specific T cell. The frequency distribution was very similar at both timepoints (Fig. 1A). Of note, 69/79 clonotypes observed at a frequency >1% were specific to another virus, mainly cytomegalovirus (CMV) and EBV. Cumulative frequencies in donors and recipients were also comparable (Fig. 1B). Most SARS-CoV-2 clonotypes were public and didn’t change in frequency at both timepoints while other public clonotypes showed a significant post-Tx increase (Fig. 1C). About 25% of clonotypes had a sequence reactive to two or more ORFs, another 25% were specific to ORF1ab and the third most frequent was against the surface glycoprotein (Fig. 1D). Generally, less than 25% of clonotypes had sequences cross-reactive to other viruses, predominantly CMV (Fig. 1E). Put together, these results suggest that unexposed individuals independently and randomly generate a large repertoire of TCRs sharing sequence homology with T cells reactive to SARS-CoV-2 (i.e., on average 5% of the total repertoire, Fig. 1B). Heterologous immunity, for instance against HCoVs, could explain some similarities in the T-cell repertoire of unexposed individuals. However, this can’t be the main driving force in more than 200 individuals, each of them being unique regarding past exposure to pathogens. Furthermore, post-Tx repertoire also strongly suggests that random somatic gene recombinations generate TCR susceptible to recognize SARS-CoV-2.

**Figure 1.**
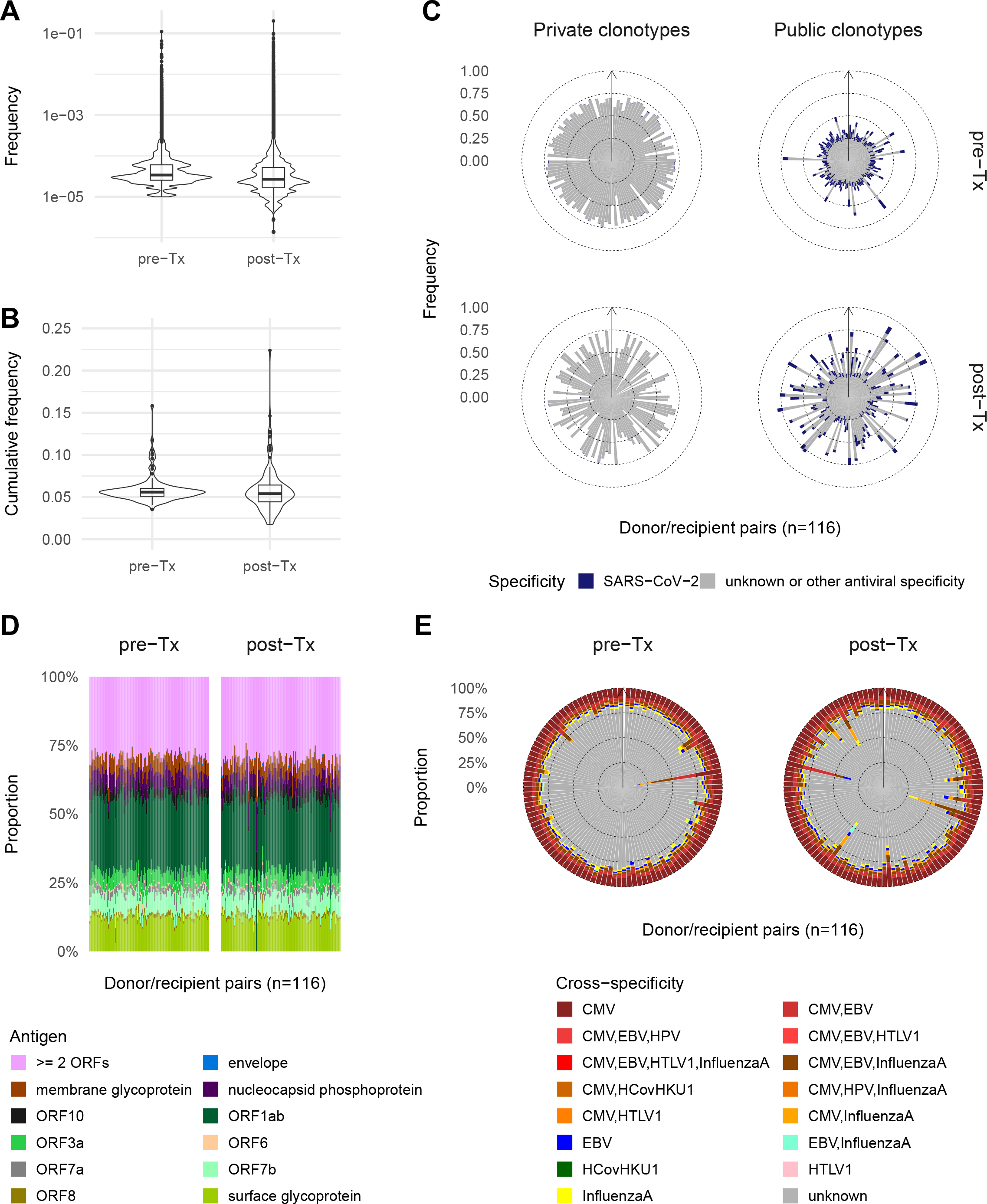
The SARS-CoV-2 TCR repertoire in 116 donor/recipient pairs. (A) frequency of SARS-CoV-2 clonotypes. (B) same as (A) but as cumulative frequency in donors and recipients one year after Tx. (C) cumulative frequency and specificity of public and private clonotypes. (D) antigenic specificity of SARS-CoV-2 clonotypes as defined according to the ImmuneCODE database, ORF: open reading frames. (E) cross-specificity of SARS-CoV-2 clonotypes against other viruses as defined according to the VDJ database.

We explored the capacity of patients at presenting SARS-CoV-2 antigenic peptides on their HLA class I molecules. The number of binders ranged from 0 to several hundreds, depending on the HLA molecule and binding level, and this was correlated to the length of the viral proteins (rho>0.95, p<1e-8, Fig. S1). Although 9-mer is the preferred length, a non-negligible number of binders are 10, 11 and 12-mer. Our data suggest that HLA-A molecules are more promiscuous at binding SARS-CoV-2 peptides, with HLA-B molecules being the most fastidious (Fig. 2A). A dual role of HLA-A and B as generalists and specialists, respectively, has been discussed ^12,13^. A significant correlation was observed between peptide overlap and HLA evolutionary divergence (HED) (mantel test, p<0.001, Fig. 2B). This is mainly driven by high divergence and low overlap at the interlocus level and smaller HED/larger overlap for some intralocus allele pairs. On average, each patient was predicted to present close to 3,000-4,000 strong and weak binders (Fig. 2C, top) with HLA-A as the prominent presenters and some interlocus overlap among presenting molecules (Fig. 2D, top). The profile was much more heterogeneous for strong binders only (Fig. 2C, bottom). The role of HLA presenters as generalists was evident, while HLA-B and HLA-C were covering much less peptides to a few exceptions. There was almost no interlocus overlap (Fig. 2D, bottom). The observed inter-individual variability, at least for strong binders, is intriguing in the light of recent studies suggesting that functional differences beyond HLA polymorphism could be related to COVID-19 outcome (e.g., susceptibility, severity and mortality) ^7,14^. Variability in peptide binding has also been described in different populations ^12^. Both qualitative and quantitative aspects of binding could affect viral infection as discussed in ^13^. Finally, we could observe that some antigenic peptides were similar between SARS-CoV-2 and HCoV strains, with 186 shared peptides and 621 and 1120 peptides differing by one or two amino acids, respectively.

**Figure 2.**
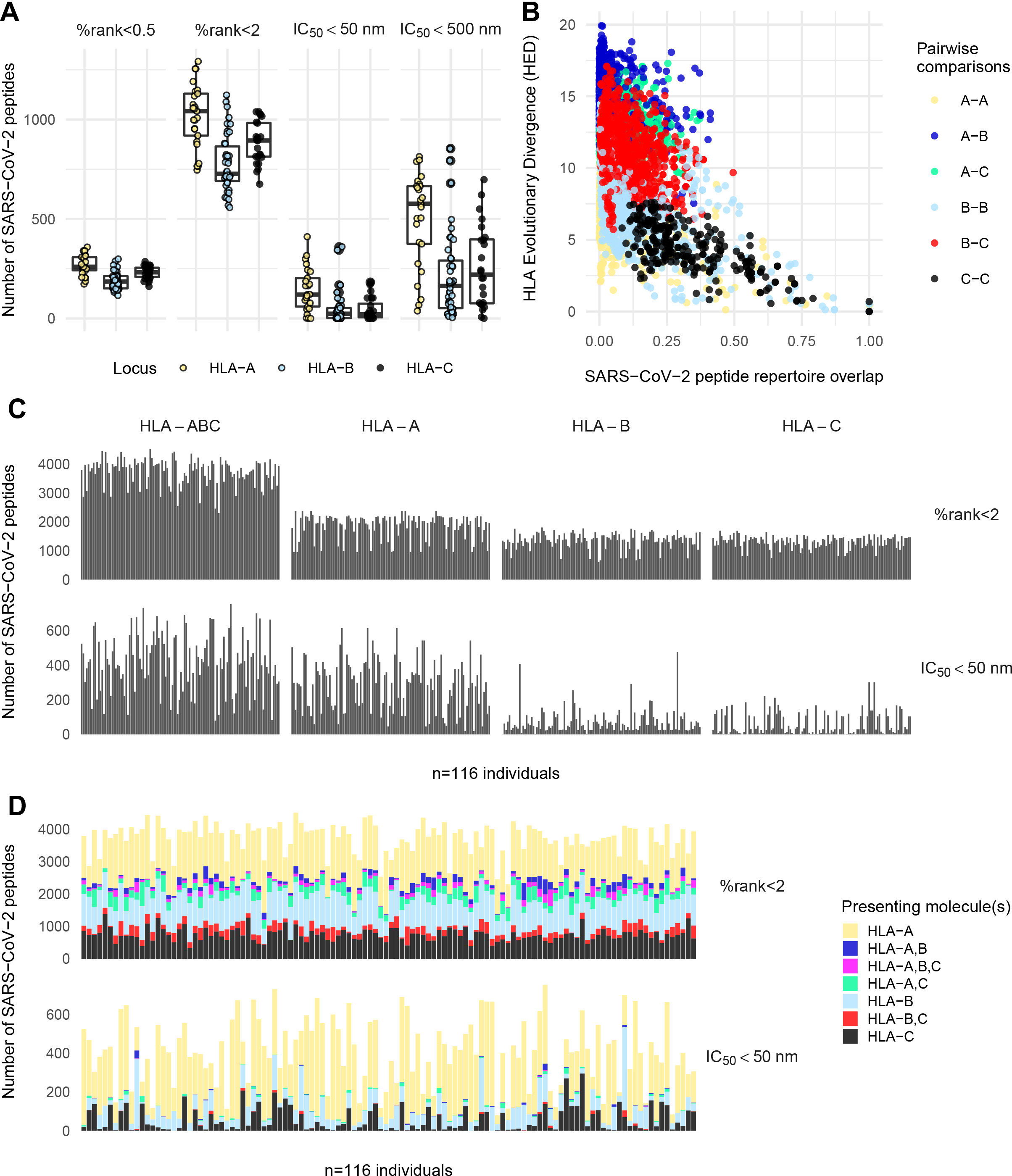
SARS-CoV-2 peptide binding predictions. (A) number of peptides presented by HLA class I alleles sorted by locus, using both prediction scores at two binding thresholds. (B) relationship between peptide overlap and HED. (C) size of the HLA class I peptide repertoire in 116 recipients for peptides considered are strong and weak binders according to percentile rank score and strong binders according to affinity score. The data are presented for combined loci and at the single locus level. (D) same as (C), but only for the combined loci and by distinguishing the presenting HLA molecule(s).

This study suggests that redundancy is the key concept behind the multifaceted and complex interactions involving TCRs and pHLAs. Each individual seems to be equipped with a diverse repertoire of T cells able to recognize a large set of SARS-Cov-2 peptides in most HLA backgrounds, however with some subtle but possibly relevant differences. Of course, this can only work in the context of a functional immune system (e.g., with controlled levels of inflammation and adequate signaling), with serious consequences in immunocompromised individuals.

## Online content

### Methods

#### Patients and donors

Patients receiving an alloHSCT graft in Geneva between 2000 and 2016, mostly as a treatment for hematological malignancies, were selected for this study. Only patients with full donor chimerism and without sign of relapse at one year were included. The cohort consisted of 116 donor/recipient pairs with characteristics described in details in ^10^. Of note, 101 pairs were 10/10 matched (i.e., for HLA-A, B, C, DRB1 and DQB1) while the 15 others carried one or sometimes more than one HLA mismatch (e.g., 4 haploidentical pairs). This dataset will allow investigating the pre-pandemic anti-SARS-Cov-2 T-cell repertoire in a large panel of healthy controls (donors) and full chimeric recipients at one year after engraftment.

#### HLA genotyping and chimerism

Ficoll purified peripheral blood mononuclear cells (PBMCs) were used to extract DNA on an automatic system (QIAGEN GmbH, Hilden, Germany). High resolution HLA typing was performed by reverse PCR-sequence-specific oligonucleotide microbead arrays and high throughput sequencing (One Lambda, Canoga Park, CA, USA) or PCR-sequence-specific primers (Genovision, Milan Analytika AG, Switzerland). Chimerism testing was performed by STR analysis (AmpFlSTR^®^ Identifiler, Invitrogen-Thermofisher, Waltham MA, USA), the detection sensitivity is <3% (i.e., patients were included if the donor chimerism was ≥97%).

#### Immunosequencing

High throughput sequencing of the TCR CDR3β region was performed on Illumina MiSeq and HiSeq platforms following a multiplex PCR of all VDJ segments according to manufacturer instructions (Adaptive Biotechnologies ImmunoSEQ© assay) ^15,16^. The two timepoints considered were just prior transplantation in the donors (pre-Tx) and at one year in recipients (post-Tx). The data were analyzed at survey resolution targeting a total of 120,000 T cells. The productive rearrangements were retrieved from the ImmunoSEQ© analyzer platform and formatted for the analyses to be carried out in R with the help of GNU/Linux scripts. Clonotypes with the same CDR3 amino acid sequence but distinct nucleotide sequences (i.e., carrying a synonymous substitution) were pooled for the analysis.

#### Antiviral specificity of clonotypes

Clonotypes in our cohort were considered as specific to SARS-CoV-2 if their CDR3β sequence exactly matched one of the 143,938 unique SARS-CoV-2 specific T-cell clone sequences available in the ImmuneCODE-MIRA-Release002.1 database (https://clients.adaptivebiotech.com/pub/covid-2020) ^17^. For each sequence listed in the database, the stimulating antigen(s) used to derive peptide pools or the ORF(s) used in transgene experiments are provided. Of note, some of the ImmuneCODE clonotypes defined by immunosequencing with the same CDR3β sequence were sometimes reactive to several stimulating antigens.

In our previous analysis ^10^, using a limited number of published CMV clones ^18^, we could show a significant increase of CMV-specific clonotypes at one year in recipients, especially in case of a documented CMV infection/reactivation but also in CMV seropositive donor/recipient pairs (i.e., D+/R+). As T cells are intrinsically cross-reactive, we investigated whether the anti-SARS-CoV-2 clonotypes retrieved in the cohort were sharing an homologous CDR3β sequence to sequences of clones known to be reactive to other viruses and to which donors and recipients could have potentially been exposed or had a clinically documented infection. For this purpose, we downloaded the sequences of antiviral clones specific to CMV (n=18,205 unique sequences in addition to the ones already available from ^18^), EBV (4,227), Influenza A (4,486), HPV (14), HSV2 (31), HTLV1 (167) and HCoV-HKU1 (48) from the VDJ repository ^19^. Clonotypes observed in the cohort and exactly matched to one of these sequences were attributed its antiviral specificity (sometimes clones having the same CDR3β sequence in VDJ were documented to be cross-reactive to several viruses). Furthermore, as the sequences in the ImmuneCODE database greatly outnumber those in VDJ, we extended the approach by considering that clonotypes in the cohort differing by at most one amino acid substitution to a given sequence in the VDJ database actually shared the same antigenic specificity. It was recently shown that clustering methods used to define the unknown antigenic specificity of a given TCR perform relatively well in this context ^20^.

All the antiviral clonotypes defined above are public, as they are observed at least once in ImmuneCODE or VDJ databases and at least once in our cohort. However, we made an additional distinction between private and public anti-SARS-CoV-2 clonotypes in the cohort according to the following definition: (1) observed only in one given donor/recipient pair or (2) at least in two pairs (either in the donor, recipient or in both), respectively.

#### HLA class I peptide binding predictions, peptide overlap and HLA evolutionary divergence

The reference proteomes of SARS-CoV-2 (UP000464024) and common cold coronaviruses HCoV-HKU1 (UP000001985), HCoV-229E (UP000006716), HCoV-HL63 (UP000167796) and HCoV-OC43 (UP000180344) were downloaded as FASTA sequences from www.uniprot.org. These data were then submitted to the NetMHCpan 4.1 server available at https://services.healthtech.dtu.dk/service.php?NetMHCpan-4.1 to perform HLA class I binding predictions ^21^. The predictions were performed on all possible 9, 10, 11 and 12-mer derived from SARS-CoV-2 and HCoVs and with each of the 91 alleles (HLA-A=25, HLA-B=42 and HLA-C=24) observed in our patients at high resolution ^22^. While 9-mer represent the canonical length of peptides bound by HLA class I molecules, peptide length preferrences vary in a molecule-dependant manner and non-9-mer dominant epitopes are well documented ^23^. NetMHCpan is trained on both peptide binding affinity experiment and mass-spectometry eluted ligand data and can predict the binding of any peptide to a given HLA molecule with good or relatively good accuracy. The binding is reported either as an affinity score (i.e., thresholds of 50nM and 500nM represent strong and weak binders, respectively) or as a percentile rank score compared with a set of natural peptides (i.e., %rank<0.5 and %rank<2 represent strong and weak binders, respectively). The rank score is expected to perform better (i.e., to provide an higher sensitivity in ligand identification at a given specificity value) than the affinity score and is usually recommended ^24^. However, this approach makes the strong assumption that every HLA molecule binds an identical number of peptides. Actually, this is not always true as some HLA molecules were shown to bind peptides at different affinity threshold and to have distinct repertoire sizes ^25^. These differences are biologically relevant ^13^, we thus considered both the percentile rank and affinity scores for a more complete overview of SARS-CoV-2 binding properties in the cohort. With the same goal in mind, the data were analyzed both at the single and multi-locus levels in order to estimate the theoretical capacity of each individual to present a set of SARS-CoV2 derived peptides for potential T-cell recognition. This analysis was limited to HLA class I alleles, as predictions are usually more accurate than for HLA class II alleles and also because constitutive HLA class II expression is restricted to a few specialized cell types. In order to avoid pseudoreplication, we only considered the HLA class I types of recipients, as all donors but eight are fully matched with their recipient at HLA-A, -B and -C and will exhibit the same binding profile.

Each HLA molecule is expected to bind a specific set of antigenic peptides (i.e., a peptide repertoire). Peptide repertoires of different HLA molecules will exhibit some degree of overlap according to the sequence similarity observed in the peptide binding region (PBR). However this relationship is not straightforward as polymorphism influences the immunopeptidome in several interlinked ways, including the repertoire of bound peptides, the quantity of peptides displayed at the cell surface and the stability and conformation of pHLA ^26^. A metric called HLA Evolutionary Divergence (HED) has been recently proposed and estimate the sequence divergence between PBRs of HLA molecules as a surrogate for their peptide binding properties ^27^. In other words, two divergent HLA alleles are expected to present a broader set of peptides for T cell recognition (i.e., less peptide overlap) than other combinations of alleles with a low HED. The HED among the 91 HLA class I alleles observed in our cohort was computed with a Perl script (https://sourceforge.net/projects/granthamdist/) and compared with the overlap of SARS-CoV-2 peptides predicted to bind to the corresponding molecules as estimated by the Jaccard similarity index.

#### Bioinformatic and statistical analyses

All the analyses were performed in R (version 4.0.3) using the RStudio integrated development environment (IDE) with the packages ggplot2, reshape2, grid, gridExtra, scales and ggpubr.

## Acknowledgements

This study was supported by the Swiss National Science Foundation (grant #310030_173237/1), the Academic Society of the University of Geneva, IRGHET (International Research Group on unrelated Hematopoietic stem cell Transplantation), the Dr Henri Dubois-Ferrière Dinu Lippatti foundation and the Philantropy Settlement.

The authors are grateful to the technicians of the LNRH for their most efficient support for HLA typing.

## Author contributions

J.V., S.F.-L., and S.B. conceptualized the study; S.B. performed the bioinformatic and statistical analysis; S.B. and J.V. drafted the manuscript; all authors assembled the data, critically reviewed and edited the manuscript, and approved the final version.

## Additional information

**Supplementary figure 1 legend**

Number of peptides presented by each of 91 HLA class I molecules observed in 116 patients. The number of peptides is shown individually for the 17 SARS-CoV-2 proteins used in the predictions and according to their length (i.e., 9, 10, 11 and 12-mer). (A) number of strong and weak binders according to the percentile rank score. (B) number of strong and weak binders according to the affinity score.

