## Supplementary figures and images for "HLA variants and TCR diversity against SARS-CoV-2 in the pre-COVID-19 era"

### Supplementary figure 1

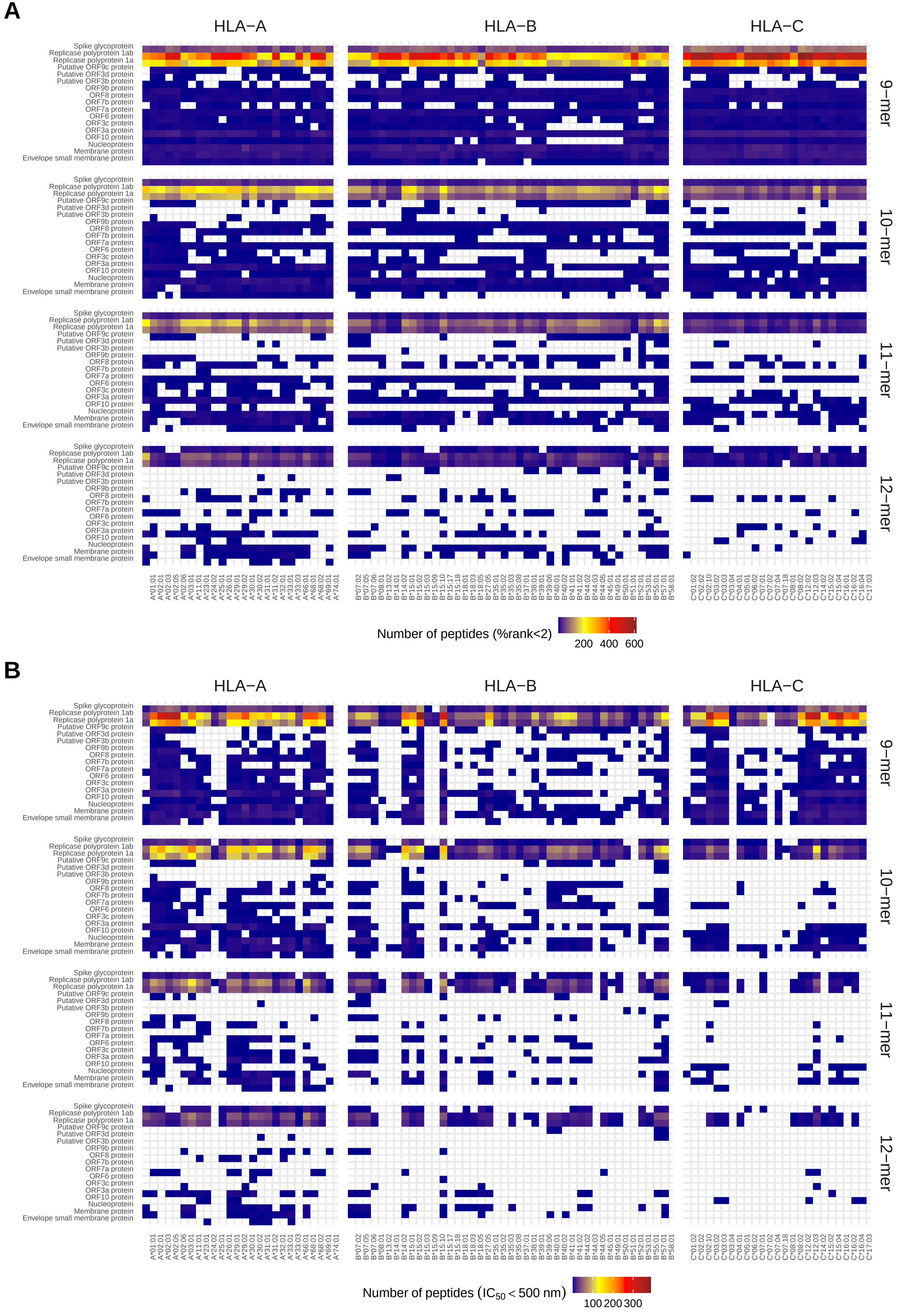
